# DREADD agonist Clozapine-N-oxide, but not Compound 21, impairs spatial memory encoding

**DOI:** 10.64898/2026.01.11.698821

**Authors:** Brayden Bunce, Anna A. VanKampen, Annie He, Sara J. Aton, Frank Raven

## Abstract

Chemogenetic studies using Designer Receptors Exclusively Activated by Designer Drugs (DREADDs) enable the precise manipulation of neuronal activity in specific brain regions and cell types. DREADDs are widely used to dissect neural circuits underlying animal behavior, including learning and memory. Clozapine-N-Oxide (CNO), a metabolite of clozapine and one of the earliest-developed ligands for muscarinic DREADDs, was initially considered pharmacologically inert. However, CNO is now known to undergo back-metabolism to clozapine, leading to undesired off-target behavioral effects, including alterations in locomotion and anxiety-related behaviors. New ligands such as Compound 21 (C21) have been developed to improve selectivity and reduce these effects. However, despite their widespread use, no studies to date have directly compared the effects of CNO and C21 themselves on specific cognitive processes such as hippocampus-dependent memory formation. Here, we measured acute effects of CNO and C21 on spatial memory encoding, using an object-location memory (OLM) paradigm in male and female mice. We also quantified encoding-associated hippocampal principal neuron and parvalbumin (PV^+^) interneuron activity by measuring cFos expression in these populations. Across dorsal hippocampal subregions, neither ligand altered overall neuronal activity nor PV^+^ interneuron activity during encoding. Nonetheless, we find that CNO administration impairs OLM encoding, while C21 does not. Together, these findings highlight a previously unrecognized behavioral effect of CNO administration on hippocampus-dependent memory formation—even in the absence of DREADD expression—and indicate that C21 may be a preferable ligand for chemogenetic studies examining memory and hippocampal function.

## INTRODUCTION

Chemogenetic tools (e.g., DREADDS) are widely used to study how specific neuronal populations contribute to behavior, learning, and memory, through selective manipulation of their activity (1, 3, 4). The commonly used excitatory (hM3Dq) and inhibitory (hM4Di) DREADDs were designed via modification to human muscarinic acetylcholine receptors to have limited response to endogenous neurotransmitters, and be activated by administration of a synthetic ligand (1, 3). Clozapine-N-oxide (CNO), a metabolite of the sedative clozapine initially assumed to be pharmacologically inert, was one of the first ligands developed to activate these muscarinic DREADDs (1). Experimental use of CNO for chemogenetic manipulations has been widely adopted, and recent studies have reported minimal or no off-target physiological or behavioral effects of CNO in animals lacking DREADD expression (5). These findings have contributed to the widespread use of CNO in behavioral studies.

However, recent rodent and primate studies have shown that CNO can be converted back into biologically active clozapine *in vivo* (6-8), leading to activation of endogenous neurotransmitter receptors (9-11). This activity may influence behavior even in the absence of DREADDs—for example, changing locomotor activity, anxiety-like behavior, and reward processing in wild-type animals (6, 7, 12, 13). Newer DREADD ligands, including Compound 21 (C21), have been designed to avoid off-target effects arising from CNO back-metabolism (14-16). Early work reported that C21 can be used as an alternative to CNO, with fewer nonspecific behavioral effects (13, 14, 17). Although several studies have compared CNO or C21 effects in wild-type animals, none have directly tested how these ligands affect hippocampal-dependent spatial memory. In the present study, we tested whether CNO or C21 alters encoding of object-location memory (OLM) in male and female mice. We quantified activity-driven protein expression, in putative principal neurons and parvalbumin-expressing (PV+) interneurons throughout the dorsal hippocampus, to determine whether any behavioral effects were accompanied by changes in encoding-associated hippocampal network activation (18, 19). By comparing CNO and C21 effects on these endpoints vs. those of vehicle treatment, we aim to identify DREADD ligand-specific effects that may influence interpretation of chemogenetic experiments relating to spatial memory processing.

## MATERIALS AND METHODS

### Animals

Male and female *SST-IRES-CRE* transgenic mice on a C57Bl/6N background (B6N.Cg-Sst^tm2.1(SST-cre)Zjh^ Jackson) at 12–21 weeks of age were used for all experiments. To identify DREADD-independent effects of ligand administration, these mice were not transduced to express DREADDs. All mice were maintained on a 12 h light/dark cycle (lights on [i.e. zeitgeber time ZT0] at 09:00 and lights off [ZT12] at 21:00) and in a constant ambient temperature (22 ± 2 °C), and were housed in standard filter-top cages with paper chip bedding material and beneficial enrichment (Enviropack nesting material). Food and water were provided *ad libitum*. All animals were group-housed with same-sex littermates and tested in the same procedure room. All housing and experimental procedures were approved by the University of Michigan Institutional Animal Care and Use Committee (IACUC; protocol PRO00011982).

### Drugs

Clozapine N-oxide (CNO; 4936, Tocris) and DREADD agonist compound 21 (C21; 5548, Tocris) were dissolved in 10% DMSO in saline, and delivered at 3 mg/kg via intraperitoneal (i.p.) injection.

### Object-location memory paradigm

The object-location memory (OLM) behavioral task was performed as described previously (20, 21). Each mouse was handled for 5 consecutive days (2 min/day) by the same experimenter as who would subsequently perform behavioral testing. During the last two days of handling, each mouse received i.p. saline injections to habituate them to injection procedures. On the last day of handling, mice were habituated to the testing arena. This arena was a rectangular 40 cm x 30 cm x 30 cm (length x width x height) chamber made from PVC, with an open top, transparent bottom, two grey walls, and two walls with a black and white vertical or horizontal line pattern as spatial cues. Each mouse was allowed to freely explore the arena for 5 min, with no objects present, during the habituation period. The following day, 30 min before OLM training (ZT 23.5), mice were injected either with CNO, C21, or vehicle (10% DMSO in saline) i.p. OLM training started at lights on (ZT0), during which mice were returned to the OLM arena containing two identical objects, placed symmetrically in the arena and equidistant from the arena walls, which they were allowed to freely explore for 10 min. OLM testing occurred 24 h following the training at lights on (ZT0). During testing, mice were presented with the same objects in the arena for 10 min, with one of the two objects in its original location, and one moved to a novel location. Specific pairs of objects and locations for moved objects were randomized and counterbalanced between treatment groups. OLM performance was quantified as a discrimination index (DI), which measures preferential interaction for the displaced object. DI = (time exploring the moved object – time exploring the stationary object) / (total exploration time). Animals exploring only one or neither of the objects during training, or exploring for less than a total of 10 s during testing, were excluded from OLM analysis. Two weeks following initial OLM training and testing, mice underwent a second round of OLM training, using a novel set of objects. 30 min prior to training, mice were injected with either CNO, C21, or vehicle, with each mouse receiving a different treatment than in the prior OLM training using a counter balanced crossover design. 90 min following training, mice were euthanized with an overdose of sodium pentobarbital, and underwent transcardial perfusion with ice cold PBS followed by 4% paraformaldehyde, in order to quantify encoding-driven expression of neuronal activity marker cFos in the dorsal hippocampus.

### Immunohistochemistry

Brains were harvested and post-fixed for 24 h, then were cut into 80 μm coronal sections using a vibratome (Leica) for immunohistochemical labeling (21, 22). Briefly, sections were washed with phosphate-buffered saline (PBS), blocked overnight in PBS with 5% normalized donkey serum (NDS) and 0.5% Triton-X100, and incubated with primary antibodies targeting cFos (Abcam, ab190289; 1:500) and parvalbumin (Millipore, MAB1572; 1:500) in PBS + 5% NDS + 0.5% Triton-X100 for 72 h. Following primary antibody incubation, sections were washed in PBS with 0.5% TX-100, incubated with secondary antibodies (goat-anti-rabbit647, Invitrogen A21245 1:750, and goat-anti-mouse488, Invitrogen A11001, 1:750) in PBS with 5% NDS and 0.5% Triton-X100 for 72 h. Following washing overnight, sections were mounted and coverslipped using ProLong Gold (Invitrogen, P36930).

### Tissue imaging and cell quantification

Following immunostaining, hippocampal subregions CA1, CA3, and dentate gyrus (DG) were imaged on a Leica SP8 confocal microscope. Acquisitions for each region, using a 10× objective, spanned a total depth of 32 μm (step size 8 μm). Identical laser exposure times and gain was used for all images of the same hippocampal subregion. Quantification of cFos^+^ cells and PV-cFos colocalization was carried out by experimenters blinded for treatment conditions, to calculate total cFos^+^ cell density and % PV^+^ interneuron cFos^+^ expression within each subregion.

### Statistical analyses

Experimenters blinded to treatment conditions performed statistical analyses using GraphPad Prism 10 (v10.2.3). For comparisons involving treatment alone, both treatment groups (CNO and C21) were compared to vehicle treatment using one-way ANOVA. For analyses including sex as an additional factor, two-way ANOVAs with treatment and sex as factors were performed. For data with non-normal distributions, the non-parametric Kruskal–Wallis test was followed by Dunn’s *post hoc* tests to compare each treatment group to vehicle. Trends were reported when p < 0.1, and statistically significant differences were reported when p < 0.05. All data are presented graphically as mean ± S.E.M.

Outlier screening was performed using the ROUT method in GraphPad Prism (Q = 1%). Values flagged by ROUT were examined for evidence of technical artifacts, including abnormally small or truncated regions of interest for immunohistochemistry data. Data points identified as technical artifacts were excluded from the corresponding analyses, while all other values were retained. Statistical results were unchanged when analyses were repeated with and without flagged values.

## RESULTS

### 3.1. Administration of DREADD agonist CNO, but not C21, impairs OLM encoding

We first examined whether CNO or C21 administered 30 min prior to OLM training altered OLM performance (**Figure 1A**). To test whether DREADD agonist administration prior to encoding impaired OLM processing, we assessed memory performance by quantifying the change in DI between OLM training and OLM testing (ΔDI). During OLM testing, one of the two objects used in training is moved to a new location, and successful OLM acquisition is associated with preferential exploration of the moved object, expressed as a positive ΔDI value. A Kruskal–Wallis test revealed a significant effect of treatment on ΔDI (**Figure 1B**; H(2) = 12.95, p = 0.0015; n = 17, 8, and 9 for Vehicle, CNO, and C21, respectively). *Post hoc* comparisons showed that ΔDI was significantly reduced in CNO-treated mice compared to vehicle-treated controls (Dunn’s test: p = 0.0012), whereas ΔDI did not differ between C21 and vehicle-treated mice (Dunn’s test: p > 0.9999). This effect did not vary between male and female mice (two-way ANOVA; main effect of treatment (F(2,28) = 6.997, p = 0.0034), main effect of sex (F(1,28) = 2.74 × 10^−5^, p = 0.9959) and treatment×sex interaction (F(2,28) = 0.2795, p = 0.7582)). Together, these results indicate that CNO treatment prior to training impairs OLM encoding, whereas C21 treatment prior to training does not.

**Figure 1.**
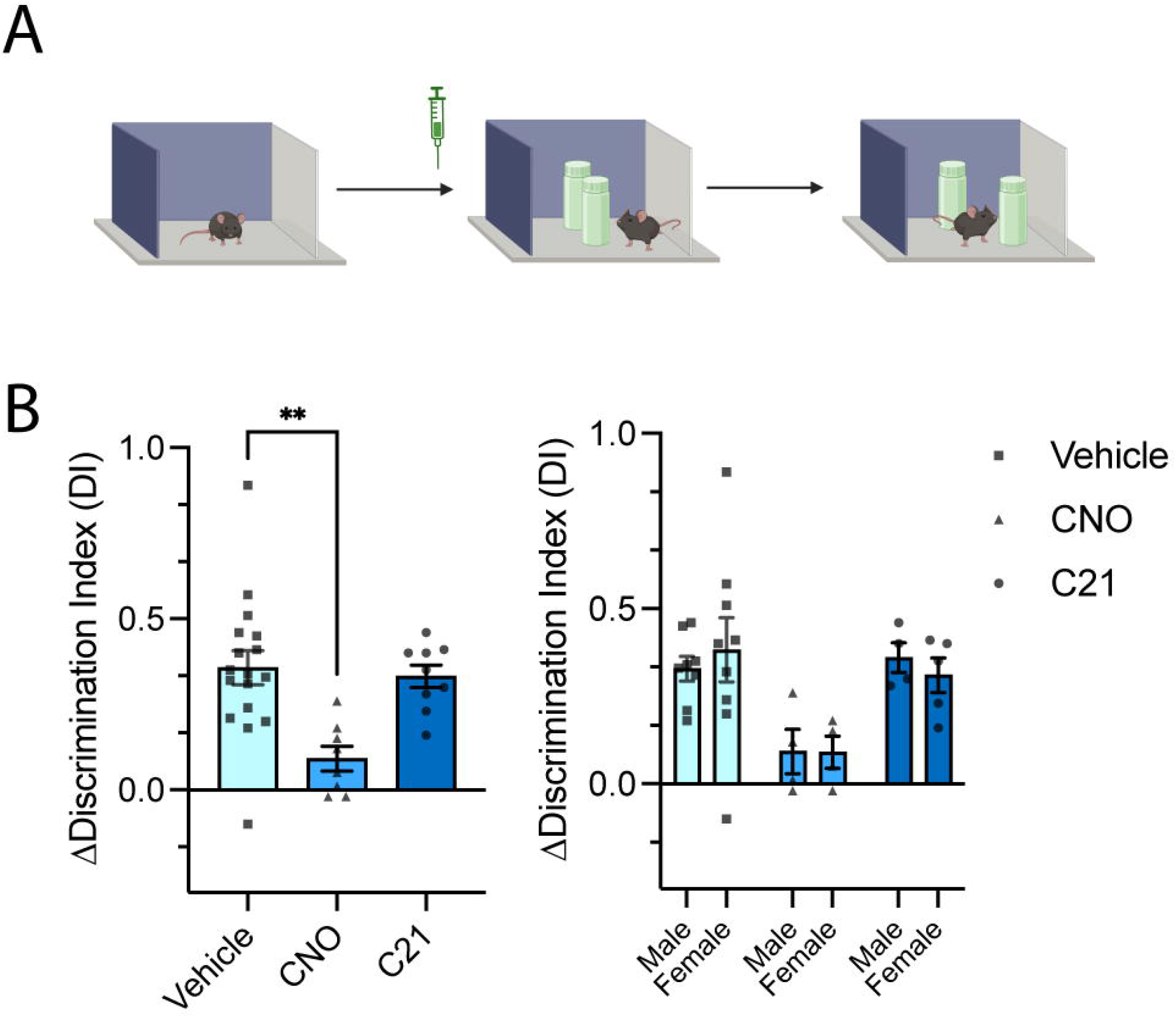
CNO, but not C21, impairs object-location memory performance. (**A)** Schematic of the object–location memory (OLM) training and testing procedure. Male and female mice (nVehicle = 17, nCNO = 8, nC21 = 9) received an injection of Vehicle, CNO, or C21 30 minutes before the training trial. Mice explored two identical objects during training, and memory was assessed 24 hours later during the test trial in which one object was displaced. **(B)** Change in discrimination index (ΔDI) across treatment groups (left) and separated by sex (right). ΔDI was significantly reduced in CNO-treated mice compared to Vehicle (Kruskal–Wallis test: H(2) = 12.95, p = 0.0015; Dunn’s *post hoc* test: p = 0.0012), whereas ΔDI in the C21 group did not differ from Vehicle (Dunn’s *post hoc* test: p > 0.9999). A two-way ANOVA including treatment and sex confirmed a significant main effect of treatment (F(2,28) = 6.997, p = 0.0034), with no effect of sex (F(1,28) = 2.74 × 10^−5^, p = 0.9959) and no interaction between treatment and sex (F(2,28) = 0.2795, p = 0.7582). Data are shown as mean ± SEM. *p < 0.05, **p < 0.01.

### 3.2 Effects of DREADD agonists on hippocampal activity patterns and PV^+^ interneuron recruitment during OLM encoding

Because the activity of the dorsal hippocampus plays a central role in OLM processing (21), we examined whether DREADD agonist treatment affected neuronal activity in this region. Brain samples were taken from mice 90 minutes after OLM training to quantify expression of cFos, which peaks around 90 min following neuronal activation (23), driven by OLM encoding in the dorsal hippocampus. Across CA1, density of cFos^+^ cells showed a trend toward a treatment effect that did not reach significance, with CNO and C21-treated mice tending to have lower cFos^+^ densities (**Figure 2A-B**; one-way ANOVA: F(2,23) = 3.070, p = 0.0658). Additional post-hoc analyses revealed that C21 significantly reduced cFos expression compared to vehicle-injected animals (Dunnett’s p = 0.0420), whereas CNO did not (Dunnett’s p = 0.5191). However, when cFos^+^ cell density was quantified within individual CA1 layers, these trends no longer existed (**Figure 2C**; one-way ANOVA for stratum oriens: F(2,23) = 0.3002, p = 0.7435; **Figure 2D;** Kruskal–Wallis test for stratum radiatum: H(2) = 1.17, p = 0.5569; **Figure 2E**; Kruskal–Wallis test for pyramidal layer: H(2) = 3.30, p = 0.1918).

**Figure 2.**
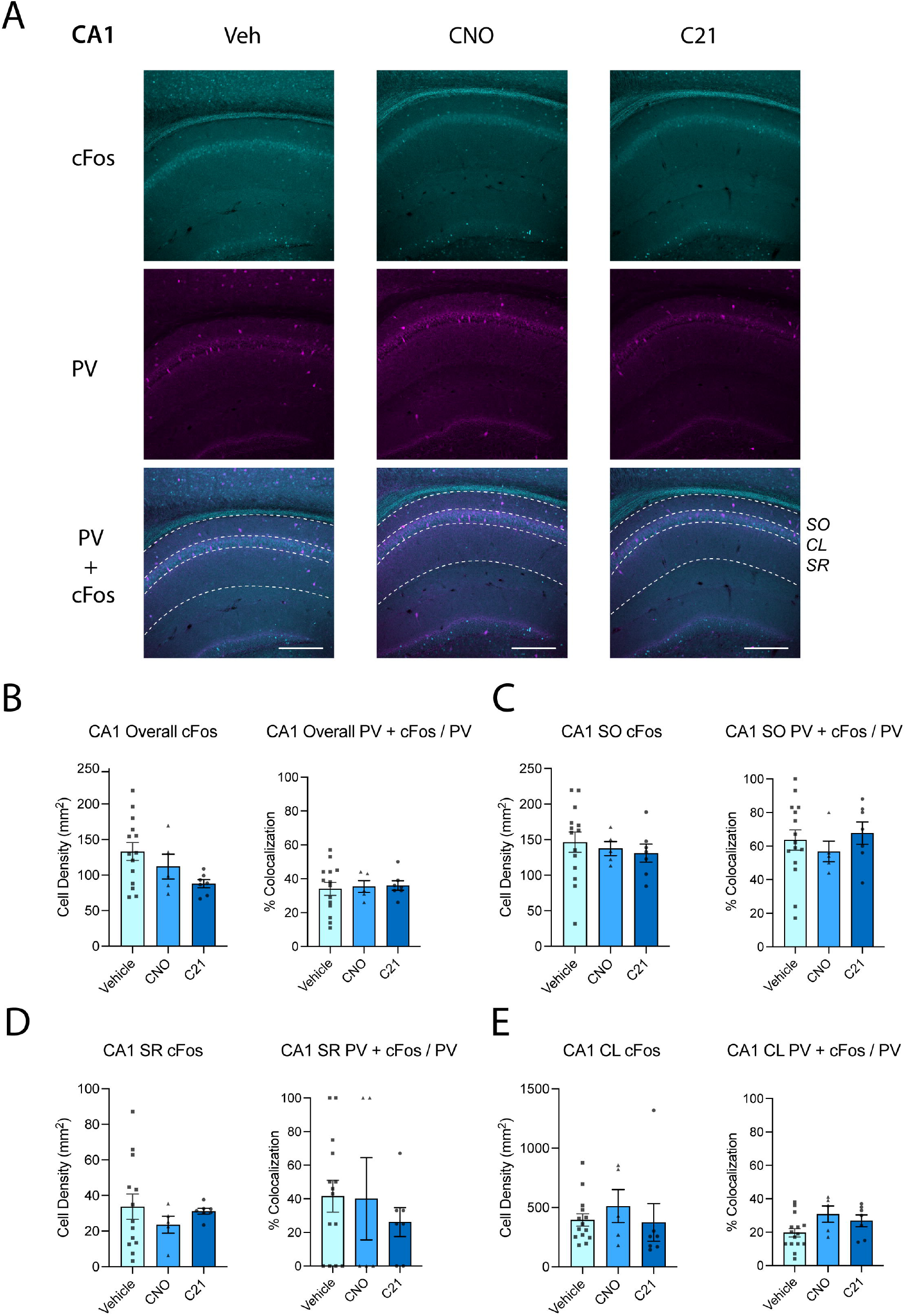
Chemogenetic treatment does not alter neuronal activity or PV^+^ interneuron activation across hippocampal subregion CA1. **(A)** Representative images showing cFos (cyan), parvalbumin (PV; magenta) immunostaining and their colocalization in dorsal hippocampal sections from Vehicle- (n = 14-15), CNO- (n = 5), and C21-treated mice (n = 7). SO: stratum oriens; CL: cell layer; SR: stratum radiatum. Scale bar = 250 μm. **(B)** Quantification of cFos^+^cell density and PV^+^ interneuron recruitment in CA1. Across all CA1 layers combined, cFos^+^ density showed a nonsignificant trend toward a treatment effect (one-way ANOVA: F(2,23) = 3.070, p = 0.0658). Quantification of PV^+^ interneuron recruitment during encoding is expressed as the percentage of PV^+^ cells that were cFos^+^ [(PV^+^+cFos^+^)/PV^+^] in CA1 overall. PV^+^ recruitment did not differ between treatment groups (one-way ANOVA: F(2,23) = 0.0685, p = 0.9340). **(C)** CA1 stratum oriens (SO): cFos^+^ density did not differ between treatment groups (one-way ANOVA: F(2,23) = 0.3002, p = 0.7435). PV^+^ interneuron recruitment did not differ between groups (one-way ANOVA: F(2,24) = 0.4065, p = 0.6705). **(D)** CA1 stratum radiatum (SR): cFos^+^ density was similar across groups (Kruskal–Wallis test: H(2) = 1.17, p = 0.5569). PV^+^ interneuron recruitment was similar across treatments (Kruskal–Wallis test: H(2) = 0.72, p = 0.6969). **(E)** CA1 pyramidal cell layer (CL): cFos^+^ density did not differ between treatment groups (Kruskal–Wallis test: H(2) = 3.30, p = 0.1918). PV^+^ interneuron recruitment remained unchanged across treatments (one-way ANOVA: F(2,23) = 2.824, p = 0.0792). Data are shown as mean ± SEM.

Because parvalbumin-expressing (PV^+^) interneurons contribute to hippocampal network dynamics underlying memory processing, we also tested whether CNO or C21 altered cFos expression among PV^+^ interneurons during OLM encoding. PV-related activity was quantified as the proportion of PV^+^ cells that were cFos^+^ [(PV^+^+cFos^+^)/PV^+^] across hippocampal subregions. In CA1 overall, PV^+^ interneuron recruitment during encoding did not differ between treatment groups (**Figure 2B**; one-way ANOVA: F(2,23) = 0.0685, p = 0.9340). PV^+^ recruitment also remained unchanged within CA1 stratum oriens (**Figure 2C**; one-way ANOVA: F(2,24) = 0.4065, p = 0.6705), stratum radiatum (**Figure 2D**; Kruskal–Wallis test: H(2) = 0.72, p = 0.6969), and the pyramidal cell layer (**Figure 2E**; one-way ANOVA: F(2,23) = 2.824, p = 0.0792).

DREADD agonist treatment also had no effect on cFos^+^ cell density in other dorsal hippocampal subregions. In CA3, cFos^+^ cell density did not differ between groups (**Figure 3A-B**; one-way ANOVA: F(2,24) = 0.0030, p = 0.9970). In the DG, neither the superior blade (**Figure 3C-D**; one-way ANOVA: F(2,24) = 0.1543, p = 0.8578) nor inferior blade (**Figure 3E**; one-way ANOVA: F(2,24) = 1.934, p = 0.1665) showed significant group differences. When the DG was analyzed as a whole (i.e., combining both blades), cFos^+^ cell density remained similar across treatments (data not shown; one-way ANOVA: F(2,24) = 0.02789, p = 0.9725). In the hilus, cFos^+^ density was also unaffected by DREADD agonist treatment (**Figure 3F**; Kruskal–Wallis test: H(2) = 4.16, p = 0.1251). Together, across all hippocampal regions measured, treatment with CNO or C21 did not alter neuronal activity levels as indexed by cFos expression.

**Figure 3.**
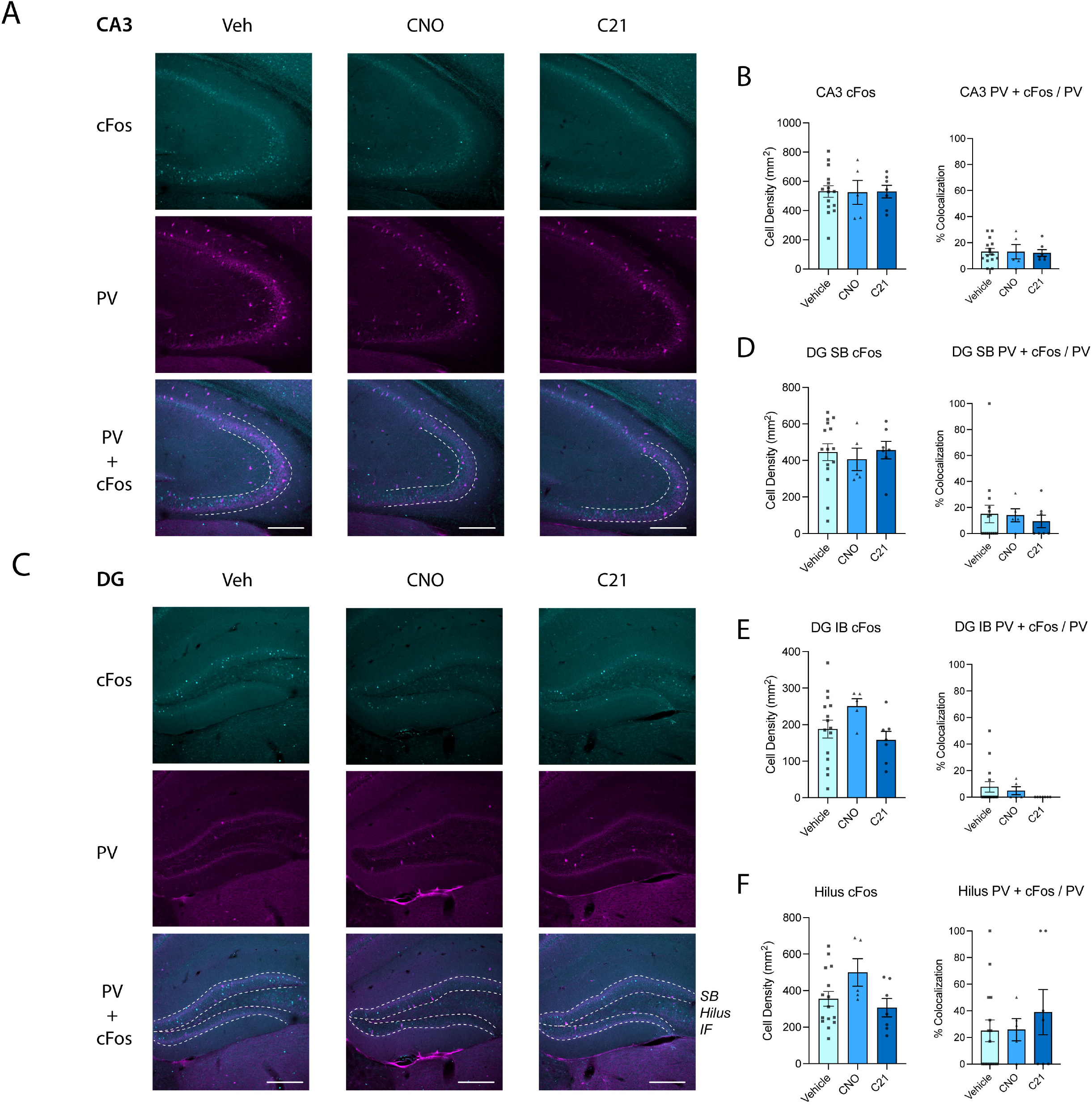
Chemogenetic treatment does not alter neuronal activity or PV^+^ interneuron recruitment during object–location memory encoding in dorsal hippocampal subregions CA3 and DG. **(A)** Representative images showing cFos (cyan), parvalbumin (PV; magenta) immunostaining and their colocalization in regions CA3 of the dorsal hippocampus from Vehicle-, CNO-, and C21-treated mice. Scale bar = 250 μm. **(B)** CA3: cFos^+^ density did not differ between groups (one-way ANOVA: F(2,24) = 0.0030, p = 0.9970). PV^+^ interneuron recruitment did not differ between treatment groups (Kruskal–Wallis test: H(2) = 0.27, p = 0.8759). **(C)** Representative images showing cFos (cyan), parvalbumin (PV; magenta) immunostaining and their colocalization in the DG of the dorsal hippocampus from Vehicle-, CNO-, and C21-treated mice. Scale bar = 250 μm. SB: superior blade; IF: inferior blade (**D**) Dentate gyrus superior blade (DG SB): cFos^+^ density was unchanged across treatments (one-way ANOVA: F(2,24) = 0.1543, p = 0.8578). PV^+^ interneuron recruitment was similar across treatments (Kruskal–Wallis test: H(2) = 0.48, p = 0.7864). **(E)** Dentate gyrus inferior blade (DG IB): cFos^+^ density did not differ between groups (one-way ANOVA: F(2,24) = 1.934, p = 0.1665). PV^+^ interneuron recruitment did not differ between groups (Kruskal–Wallis test: H(2) = 2.66, p = 0.2645). **(F)** Hilus: cFos^+^ density showed no significant group differences (Kruskal–Wallis test: H(2) = 4.16, p = 0.1251). PV^+^ interneuron recruitment remained unchanged across treatments (Kruskal–Wallis test: H(2) = 0.55, p = 0.7584). For all figure panels; Vehicle n = 15; CNO n = 5; C21 n = 7. Data are shown as mean ± SEM.

PV^+^ interneuron recruitment was likewise unchanged in these subregions of the dorsal hippocampus, including CA3 (**Figure 3A-B**; Kruskal–Wallis test: H(2) = 0.27, p = 0.8759), the dentate gyrus superior blade (**Figure 3C-D**; Kruskal–Wallis test: H(2) = 0.48, p = 0.7864), the dentate gyrus inferior blade (**Figure 3E**; Kruskal–Wallis test: H(2) = 2.66, p = 0.2645), and the hilus (**Figure 3F**; Kruskal–Wallis test: H(2) = 0.55, p = 0.7584). Together, these results indicate that neither CNO nor C21 produced detectable changes in PV^+^ interneuron recruitment during memory encoding at the time point assessed.

## DISCUSSION

Recent work has highlighted off-target behavioral effects of chemogenetic ligands, particularly CNO. Behavioral effects of CNO have been attributed back-metabolism to clozapine and/or engagement of endogenous GPCRs (6, 7, 12, 13). However, only a handful of studies have directly compared cognitive effects of CNO to those of newer ligands such as C21 (13, 14, 17). In the present study, we tested whether acute administration of CNO or C21 alters object-location memory (OLM) encoding in male and female mice. We administered each ligand before training and subsequently quantified both memory performance and histological markers of neuronal activity across the hippocampus. We found that in both male and female mice, CNO impaired memory performance, whereas C21 did not. Because CNO impaired memory encoding, we next asked whether this deficit was associated with altered hippocampal activity during memory encoding. The hippocampus is highly sensitive to neuromodulatory influences, and even subtle alterations in network state can influence memory encoding processes (24, 25). To assess whether either ligand produced broad shifts in hippocampal activity during encoding, we quantified cFos expression across hippocampal subregions at a time point when cFos levels peak following neural learning. Given the established role of parvalbumin-expressing (PV^+^) interneurons in regulating hippocampal network dynamics relevant to memory processing (17, 26, 27), we also assessed PV^+^ interneuron recruitment during encoding as PV/cFos colocalization (18). To our surprise, across all hippocampal regions examined, neither CNO nor C21 significantly changed encoding-related cFos expression, suggesting that the observed behavioral effects were not associated with altered hippocampal activity. Together, these findings indicate that CNO impairs object-location memory encoding without significantly affecting hippocampal activation or PV^+^ interneuron recruitment.

One possible explanation of this set of results is that CNO’s effects on spatial memory arise through mechanisms not captured by measurement at this single time point. While previous work from our lab has demonstrated that CNO administration does not affect contextual fear memory (18), it is possible that consolidation of OLM is impacted by CNO administration. Another possibility is that CNO, or its conversion to clozapine, subtly alters neuronal excitability, neuromodulatory tone, or network coordination in ways that impair encoding without producing large changes in hippocampal immediate early gene expression. In addition, CNO may influence neural circuits outside the hippocampus that are involved in object-location association, or in circuits regulating arousal, attention, or motivation—all of which could indirectly impact OLM encoding. Addressing these possibilities will require time-resolved approaches that assess neural activity across multiple post-training time points.

Our findings demonstrate that CNO can impair hippocampus-dependent memory encoding even in the absence of DREADD expression, extending prior reports of off-target behavioral effects associated with this ligand. Importantly, this impairment occurred without detectable changes in hippocampal activity or PV^+^ interneuron recruitment during encoding, indicating that CNO’s behavioral effects are likely mediated by more subtle or temporally specific mechanisms. In contrast, C21 did not affect memory performance or hippocampal activity measures under the same conditions, supporting its use as a more precise chemogenetic ligand. More broadly, this work underscores the importance of evaluating ligand-specific effects when interpreting chemogenetic manipulations and highlights the need for careful selection and validation of DREADD actuators for studies of learning and memory.

## ABBREVIATIONS

C21: Compound 21
CNO: Clozapine-N-Oxide
DG: Dentate gyrus
DREADDs: Designer Receptors Exclusively Activated by Designer Drugs
NDS: Normalized donkey serum
OLM: Object-location memory
PBS: Phosphate-buffered saline
PV: Parvalbumin

## ACKNOWLEDGMENTS

**Figure 1A** schematic was created using Biorender. These studies were supported by NIH R01NS118440, NIH R01MH135565, a Chan Zuckerberg Initiative Collaborative Pairs grant, and a University of Michigan Biosciences Initiative grant to SJA, and by NIA K99AG088300 and Michigan Alzheimer’s Disease Research Center grant to FR.

